# A statistical test on single-cell data reveals widespread recurrent mutations in tumor evolution

**DOI:** 10.1101/094722

**Authors:** Jack Kuipers, Katharina Jahn, Benjamin J. Raphael, Niko Beerenwinkel

**Author notes:** equal contributors (Jack Kuipers, Katharina Jahn).

## Abstract

The infinite sites assumption, which states that every genomic position mutates at most once over the lifetime of a tumor, is central to current approaches for reconstructing mutation histories of tumors, but has never been tested explicitly. We developed a rigorous statistical framework to test the assumption with single-cell sequencing data. The framework accounts for the high noise and contamination present in such data. We found strong evidence for recurrent mutations at the same site in 8 out of 9 single-cell sequencing datasets from human tumors. Six cases involved the loss of earlier mutations, five of which occurred at sites unaffected by large scale genomic deletions. Two cases exhibited parallel mutation, including the dataset with the strongest evidence of recurrence. Our results refute the general validity of the infinite sites assumption and indicate that more complex models are needed to adequately quantify intra-tumor heterogeneity.

---

The presence of mutational heterogeneity within tumors due to somatic cell evolution is known to be a major cause of treatment failure ^1,2^. With the emergence of next-generation sequencing techniques it is possible to systematically analyze individual tumors at a genetic level from admixed cell samples, and more recently from sequencing the DNA of individual tumor cells ^3,4^. These technical advances, together with a prospect of high-precision cancer therapies, have spurred the development of a variety of computational approaches to reconstruct not only the clonal structure but also the entire mutation history of individual tumors ^5-19^. A common feature of all these approaches is the use of the infinite sites assumption (ISA) ^20^ to exclude the possibility of the same genomic site being affected by multiple mutations throughout the lifetime of a tumor. However, the ISA has never been explicitly tested in the context of tumor evolution on sequencing tumor data. Only in the context of copy number alterations it has been recently suggested to allow multiple changes of the same site while still excluding recurrences of the same state ^21,22^.

The ISA is convenient to make, as it substantially restricts the search space of possible mutation histories ^23^, but its validity is unproven and hard to test, as many factors such as mutation rate, cell division rate, copy number changes and the presence of mutational hotspots influence the probability of multiple mutations hitting the same site. On larger scales, multiple mutations have been observed to affect the same gene at different genomic sites in different spatial areas and phylogenetic branches of tumors ^24,25^ indicating convergent evolution for these driver genes. Structurally different copy number alterations have also been observed to affect the same genes in ovarian cancer ^22^. This raises the specter of recurrence at the scale of individual bases and violations of the ISA (Figure 1).

**Figure 1:**
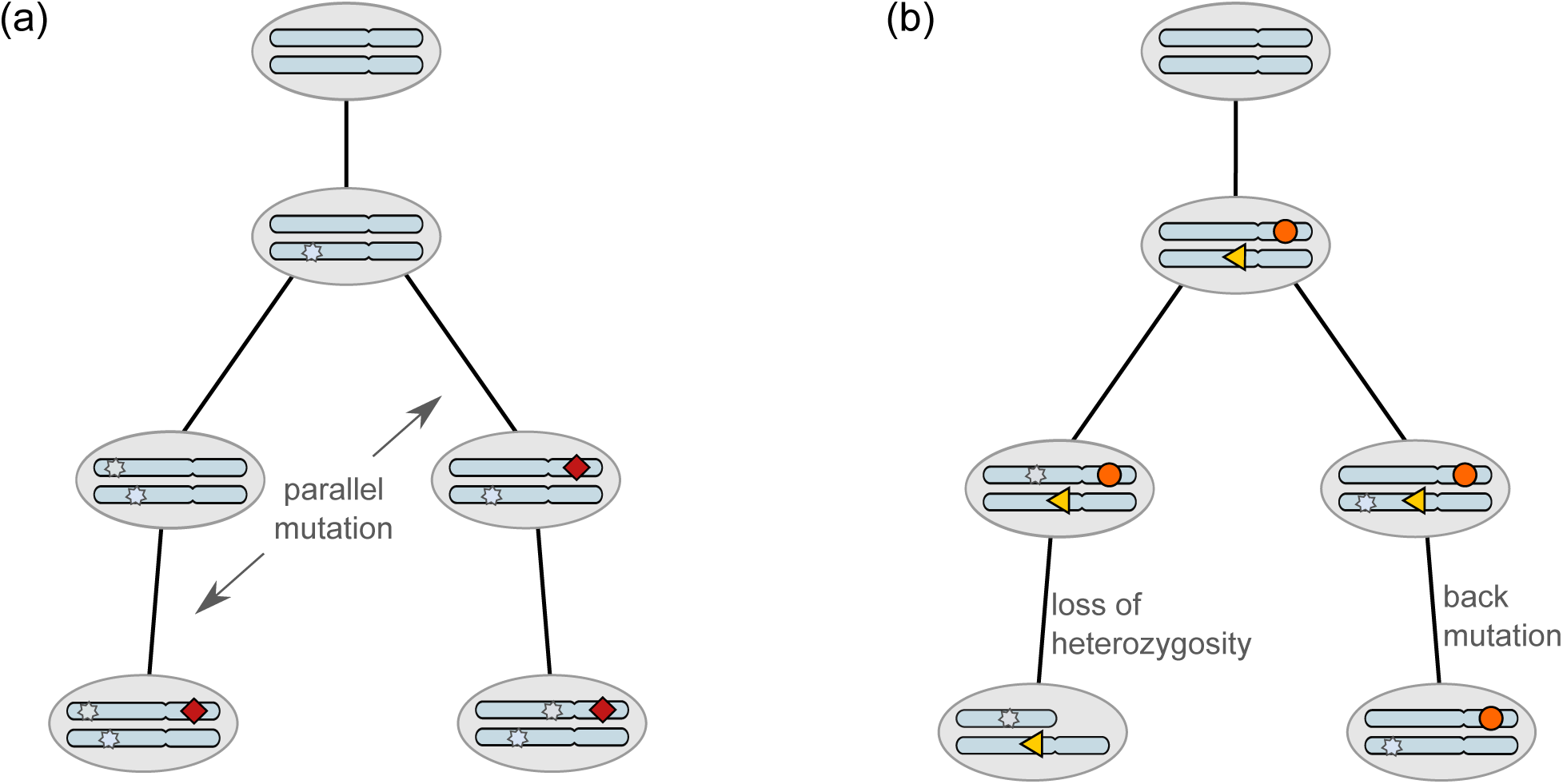
Somatic mutations occurring during tumor evolution could violate the infinite sites assumption. (a) The mutation indicated by the red diamond occurs in *parallel* in two different lineages. (b) The mutation depicted by the orange circle is lost in the left branch due to a *loss of heterozygosity*. The mutation drawn as a yellow triangle is lost in the right branch by reverting to its original state, denoted a *back* mutation.

In fact, the idea that every genomic position mutates at most once over the life-time of a tumor can be disproved by a generalization of the birthday problem (Online Methods). This is a classic math puzzle that asks for the probability that two people in a group share the same birthday. Perhaps surprisingly, this probability is already greater than 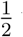 with only 23 people. Using the same reasoning and estimates of the cumulative number of stem cell divisions ^26^ and mutation rates ^27^, we found that the probability of violating the ISA in any tissue is almost 1 (Supplementary Note).

It is a different question, however, whether recurrent mutations are likely to be observed in practice, as only a small fraction of the evolutionary history is reconstructable from the limited number of tumor cells that are typically sequenced (Supplementary Figure 1). So although it is almost certain that the ISA is violated within the tumor tissue, there may still be a low chance that a violation occurs with a small set of mutations observed in a small sample of cells (Supplementary Note).

Therefore we developed a statistical framework (Figure 2) based on real tumor data to test the infinite sites model (ISM), *𝓜*_I_, that comprises all histories with a single event for every mutated site, against a model *𝓜*_F_ that allows multiple mutations at the same site, referred to as finite sites model (FSM) (Online Methods). The test is defined as a model selection problem where we compute the Bayes factor (BF) ^28,29^ of the two alternative models based on single-cell sequencing data, *𝐷*,

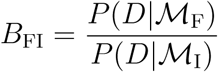

**Figure 2:**
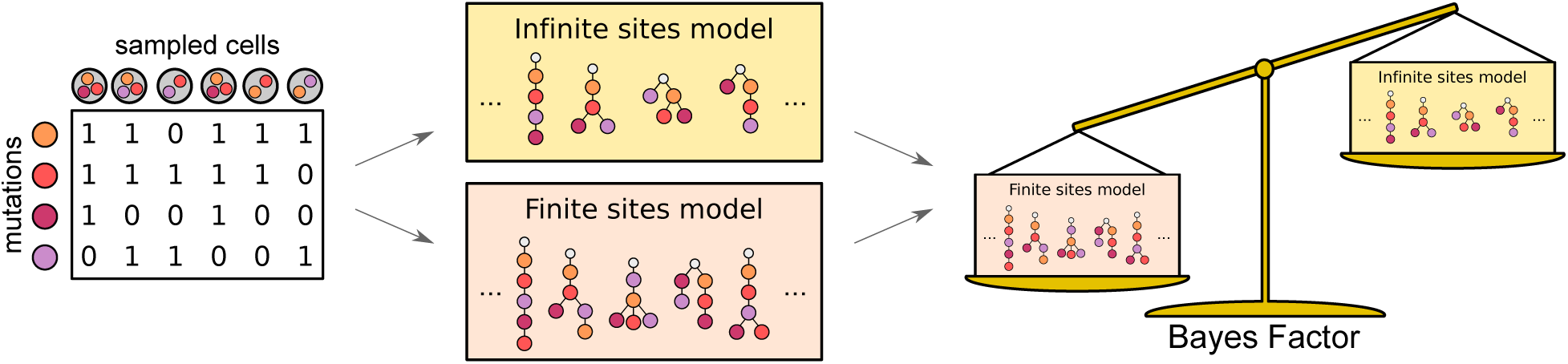
Testing the infinite sites assumption starts from the single-cell mutation data. The data is examined under both the infinite sites model of all trees with no recurrent mutations as well as under the finite sites model of trees with one recurrence. The two competing models of tumour evolution are compared on how well they explain the single-cell data, with one model selected via the Bayes factor.

When the FSM fits the data better than the ISM, the BF is greater 1, and the larger the value, the stronger is the evidence against the ISA. Neatly, the BF can be combined with estimates of the prior odds of each model to provide the posterior odds:

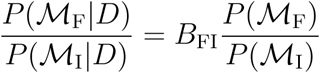

The computation of the BF is based on our earlier work on reconstructing mutation histories from mutation profiles of single cells ^18^, which we generalize here to allow a single recurrent mutation (Online Methods). The recurrent mutation can be either a back mutation, if the second event occurs in the same cell lineage, or a parallel mutation that occurs in a different lineage (Figure 1). The reconstruction accounts for the noise in single-cell sequencing data, particularly the high levels of allelic dropout.

Single-cell sequencing data can additionally be contaminated by doublets, the inadvertent sequencing of more than one cell together, with some platforms having rates as high as 40% ^30^. We observed that high doublet contamination rates affect the quality of the reconstructed mutation histories and thereby can confound the model selection process. Therefore we extended both models to account for doublets and to learn their incidence rates from the data (Online Methods).

## Results

Evaluation of our framework on simulated data sets with realistic noise levels and contamination with doublets revealed that our test has a high specificity of 95% using a BF cutoff of 1 (Supplementary Note). The sensitivity increases with the number of sequenced cells. With 2–3 cells per mutation, we find a moderate sensitivity of 50–60% with the same BF cutoff. While this means that some recurrent mutations will be overlooked, any signaling of violations of the infinite sites assumption in real data can be trusted. We analyzed nine published single-cell tumor datasets, three from whole-exome sequencing (Table 1) and six from targeted sequencing (Table 2). The details of the inferred parameters and trees are discussed in the Supplementary Note, with the results presented here.

**Table 1:**
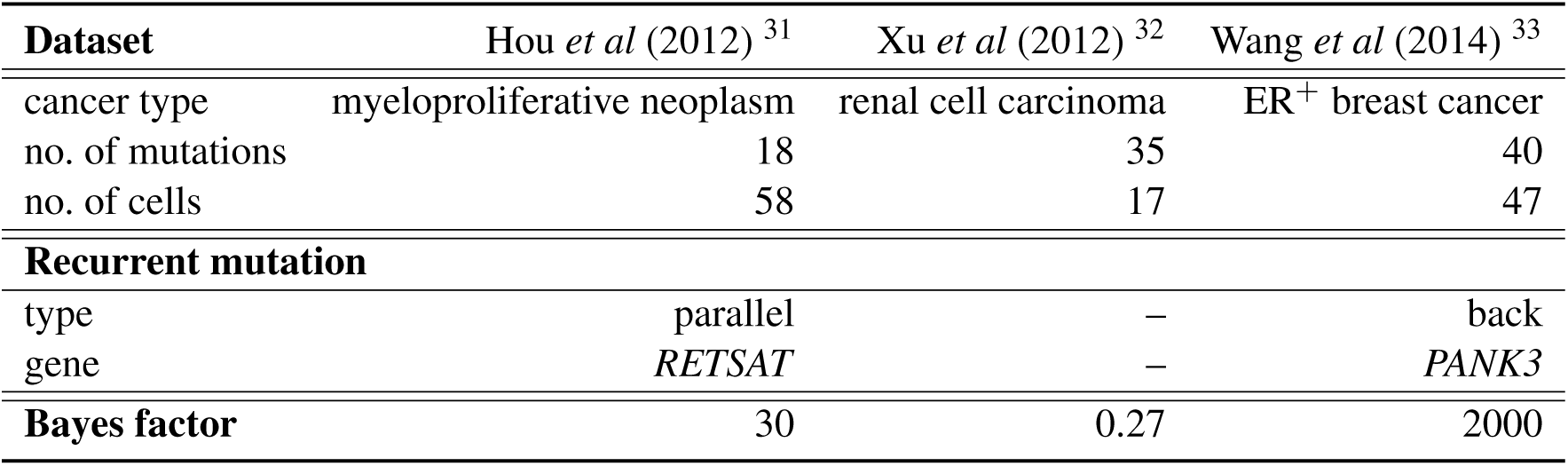
Characteristics of the three exome sequencing datasets^31-33^ along with the inferred recurrent mutations and Bayes factors.

| **Dataset** | Hou *et al* (2012) <sup>31</sup> | Xu *et al* (2012) <sup>32</sup> | Wang *et al* (2014) <sup>33</sup> |
| --- | --- | --- | --- |
| cancer type | myeloproliferative neoplasm | renal cell carcinoma | ER<sup>+</sup> breast cancer |
| no. of mutations | 18 | 35 | 40 |
| no. of cells | 58 | 17 | 47 |
| **Recurrent mutation** |  |  |  |
| type | parallel | – | back |
| gene | *RETSAT* | – | *PANK3* |
| **Bayes factor** | 30 | 0.27 | 2000 |

**Table 2:** Characteristics of the panel sequencing datasets of six leukemia patient samples^34^ along with their inferred recurrent mutations and Bayes factors. The genomic positions are according to the hg19 assembly.

| **Patient** | **1** | **2** | **3** | **4** | **5** | **6** |
| --- | --- | --- | --- | --- | --- | --- |
| panel size | 98 | 82 | 196 | 155 | 248 | 85 |
| no. of mutations | 20 | 16 | 49 | 78 | 105 | 10 |
| no. of cells | 111 | 115 | 150 | 143 | 96 | 146 |
| **Recurrent mutation** |  |  |  |  |  |  |
| type | back | back | back | back | parallel | back |
| gene | *MAL2* | *RIMS2* | *CUL3* | *IKBKB* | *C1orf105* | *SUSD2* |
| chromosome | chr8 | chr8 | chr2 | chr8 | chr1 | chr22 |
| genomic position | 120255800 | 105025789 | 225335303 | 42162708 | 172434428 | 24582021 |
| substitution | $C \rightarrow G$ | $T \rightarrow G$ | $G \rightarrow C$ | $C \rightarrow G$ | $C \rightarrow A$ | $C \rightarrow T$ |
| **Bayes factor** | $8.6 \times 10^5$ | 330 | $4.1 \times 10^{10}$ | $1.8 \times 10^7$ | $4.8 \times 10^{15}$ | $9.7 \times 10^{13}$ |

### Evidence for recurrent mutations in single-cell exome sequencing data

Looking at a *JAK2*-negative myeloproliferative neoplasm (essential thrombocythemia) for which the exomes of 58 tumor cells were sequenced, we focussed on the 18 mutations classified as cancer-related ^31^ and find evidence for a recurrence of the same point mutation in the *RETSAT* gene (Supplementary Figure 11). Both mutations are late events that have happened at the end of two neighboring branches. This recurrence is supported by a BF estimate of 30 constitutes strong evidence for a violation of the ISA.

Next, we analyzed a clear cell renal cell carcinoma for which exome sequencing data of a total of 17 tumor cells is available ^32^. Performing the model comparison based on the 35 sites informative for mutation tree reconstruction, we obtain a BF below 1. There is therefore no evidence for a violation of the ISA, although any such violation would be hard to detect with the low number of sequenced cells.

In a dataset of 47 cells of an estrogen-receptor positive (ER^+^) breast cancer with 40 informative mutation sites^33^, we found that the tree topology under both models consists of a linear chain of mutations on top of a rather branched structure further down. Under the FSM a back mutation of the early *PANK3* mutation, changes the upper tree structure substantially compared to the tree under the infinite sites model where the mutation is forced into a side branch (Supplementary Figure 13). Computing the BF, we find a value of 2000 providing very strong evidence that the model with the back mutation fits the data much better than the infinite sites model.

For the small number of cells sequenced, and assuming a uniform distribution of mutations with no selection and that all mutations are observed, we obtain the conservative estimate of the probability of the same site among 40 changing twice via point mutations to be rather small at 2.5 × 10^−5^ (Supplementary Table 5). We therefore tested loss of heterozygosity (LOH) as an alternative explanation for the back mutation: If the only allele carrying the mutation is lost at some point in the tree, sequencing descendant cells will only yield reads from the normal allele thereby mimicking a back mutation (Figure 1). Based on copy number data from breast cancer samples from the TCGA Research Network (cancergenome.nih.gov/), LOH on the *PANK3* gene occurred with a probability of approximately 2 × 10^−3^ and thereby much higher than for the uniform reversion of a point mutation among 40. Copy number estimates are also provided ^33^ for the sequenced cells, although it is difficult to determine whether LOH has occurred in the respective region. The reason being that *PANK3* is located on chromosome 5 which was amplified early in the tumor evolution. Of the sequenced cells most of them seem to still exhibit an amplification of chromosome 5, but this is less certain for all cells. Some cells may then have lost a copy later, giving a possible explanation of our observation of the back-mutation.

### Evidence for recurrent mutations in single-cell panel data

We found the strongest evidence against the ISA in single-cell sequencing data from the personalized panels of six childhood acute lymphoblastic leukemia (ALL) patients ^34^. Our test returns extremely high BFs in the range of 10^5^ to 10^15^ (Table 2) for five of the cases, and a more modest but still highly significant BF estimate of 330 for one patient sample (patient 2). For all samples apart from patient 5, the recurrent mutation is a back mutation. Looking at the trees (Supplementary Figures 14–19) we notice that for three patients the lost mutation is actually the first one that happened in their trees: They affect the *MAL2* gene in patient 1, *RIMS2* in patient 2 and *SUSD2* in patient 6. For patient 4, the lost mutation was in *IKBKB* which was also acquired in the tree trunk, while the last case, patient 3, lost a mutation in *CUL3* that was acquired further down in a branch of the tree. Interestingly also three out of the five back mutations occur on chromosome 8. The overrepresentation of reversions of early clonal mutations could hint at changing selective pressure that renders an early trunk mutation expendable or even hindering in later tumor stages. Signs of this biological possibility have recently been observed for Barrett’s oesophagus^35^.

Since LOH events are the most likely causes of back mutations, we compared to the 16 LOH events (>10kb) detected from the bulk data of the 6 leukemia patients^34^. However, the single-cell data showed that the large majority (13 out of 16) appeared in all clones and were ancestral^34^. None of the five back mutations we identified appeared in any of the LOH regions of the respective patient, emphasizing that they are unlikely to be the result of large scale deletions. smaller scale deletions. The data then indicate either smaller scale deletions or genuine back mutations with a reversion of the individual loci.

We further examined whether the LOH at the loci we identified are common in ALL. To obtain such statistics, we performed a comparison with large scale (>5Mb) copy number deletions found in a large study of 142 children and 123 adults with ALL ^36^. Patients 1 and 2 had back mutations on the 8q chromosome, which were not observed in any of the study samples, except a whole loss of chromosome 8 in one adult. Patient 3 had a back mutation at chromosome 2q, which also was not observed in any of the study samples. The back mutations for patients 1–3 therefore also do not seem to match common large-scale deletion events, but could be the result of smaller losses. The 8q back-mutations of patient 1 and 2 are close to *MYC* which plays an important role in ALL ^37^. Patient 4’s back mutation was at 8p which was lost in 4 children and 4 adults, while patient 6’s back mutation was at 22q and chromosome 22 was subject to large-scale deletion for 4 children and one adult in the ALL sample ^36^. Their large BFs could be related to these relatively common LOH events. Patient 6’s back mutation also happened to be near *IGL* which is rarely translocated with *MYC*.

For patient 5, we observed (Figure 3) a parallel mutation in *C1orf105* with a BF of 4.8 × 10^15^ so that allowing the mutation to occur twice explains the data much better than enforcing the ISA. Since sequencing bias is an unlikely explanation for the extreme BF based on analyzing the read counts in the cells (Supplementary Note), our conclusion is that we are observing here a real signal of the same genomic position mutating twice in different subpopulations of a tumor.

**Figure 3:**
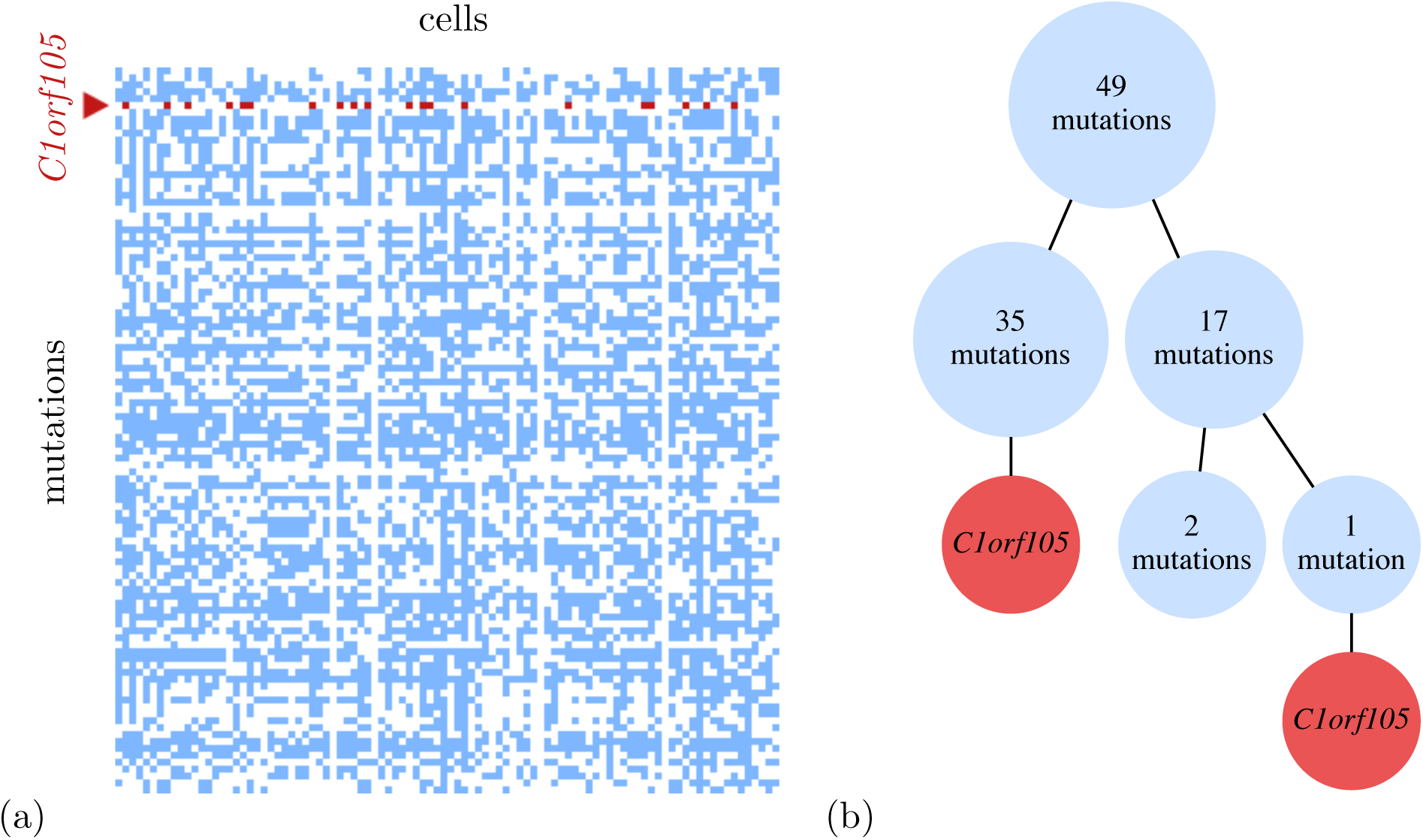
(a) The data matrix of the 105 mutations detected in the 96 single cells of patient 5 of the leukemia dataset ^34^. Unmutated positions are left white, mutations are colored blue and the recurrent mutation in *C1orf105* colored red. (b) The inferred mutational history under the finite sites model when allowing a recurrence of the point mutation in *C1orf105.* The two occurrences appear at the ends of different lineages in the tree, separated in the two branches by 35 and 18 other mutations. The very large Bayes factor of 4.8 × 10^15^ shows that allowing the parallel mutation fits the data much better than enforcing the infinite sites assumption.

### Signs of secondary parallel mutations

Since back mutations violate the ISA but may have a simpler biological cause from LOH than the single genomic position reverting, we wished to examine parallel mutations more closely because these act at the level of individual bases. In particular, we restricted our search to consider only the highest scoring parallel mutation for each dataset. This may reveal additional violations of the ISA.

For the exome data, the recurrent mutation uncovered from the myeloproliferative neoplasm ^31^ is already parallel and no other parallel mutation scored highly. No evidence for infinite sites violations was discovered for the kidney cancer ^32^, and for the breast cancer samples ^33^ no parallel mutation scored highly. For the panel data ^34^ on the other hand, we find parallel mutations for patients 1–4 with BFs larger than 1 (Supplementary Table 3). Three of them have moderate BFs, but for patient 3 we find a large BF of 2.4 × 10^6^ which indicates multiple violations of the infinite sites hypothesis.

For patient 5 we also found multiple parallel mutations. The top-scoring recurrence was already a parallel mutation (Table 2), but the second highest scoring recurrence is also parallel with a very large BF of 4.1 × 10^10^. That mutation occurs on chromosome 9 at position 139923258 (hg19) which is at the ends of the *ABCA2* and *C9orf139* genes.

## Discussion

We have developed a statistical framework to test the infinite sites assumption in single-cell sequencing data. Application of our framework to published patient data (one myeloproliferative neoplasm^31^, one renal cell carcinoma^32^, one breast tumor^33^, and six leukemia patients^34^) suggests that the assumption is frequently violated. We showed that these findings can not be explained by the background mutation rate alone, as the prior probability of mutating the same base twice among a selected set of bases is low if mutations are spread uniformly across the genome (Supplementary Table 5).

Most of the observed violations of the infinite sites assumption present as back mutations, typically as the loss of an early clonal mutation. This may be the result of random losses of passenger mutations, but observing this pattern in many patient samples would also be compatible with selection driven by the micro-environmental or the genetic context. For example early driver mutations may become obsolete once the tumor is established, or may even hinder the tumor at later stages so their loss becomes positively selected for. Hints of changing selective pressures on particular aberrations have recently been observed for Barrett’s oesophagus^35^. Loss of a copy of the p16-locus seemed to provide a fitness advantage for clones experiencing acid reflux but a disadvantage when the acid is suppressed under treatment. Clones that regain the p16 copy could then potentially experience positive selection. For half of the leukemia patients the backmutation occurs on chromosome 8 pointing to a particular role in the development of the disease. A simpler explanation for back mutations is LOH, the loss of a chromosomal segment that comprises a mutated site. In tumors rich in copy number alterations such an event would have a reasonably high prior probability, as the same site is much easier hit by two or more such large-scale alterations than by two point mutations. In the leukemia dataset ^34^, the back mutations we identified did not occur in genomic regions affected by large scale deletions. While our findings on the incidence of back mutations are limited to the small number of patient samples available at this point, they may be of importance in the context of treatment strategies that target early trunk mutations in cancer therapy. Our method can be used to generate the trunk mutations more accurately, as evident particularly for the breast cancer sample ^33^ (Supplementary Figure 13).

We also found evidence for parallel mutations in two of the studied cases, patient 5 of the leukemia dataset ^34^ (Figure 3) and the *JAK2*-negative myeloproliferative neoplasm ^31^. In both cases, the two mutation copies appear at the end of different lineages, which could also point to selective pressure from the tumor environment or the genetic context. Having corrected for the possibility of doublet samples in our model, the event of a mutation hitting the same site twice appears here to be the most plausible explanation. Conservative estimates of the prior odds of recurrent mutations among a small set of mutations of interest were obtained by spreading mutations uniformly across the genome and assuming that all mutations are observed (Supplementary Table 5). With these low prior estimates, the posterior probability of the infinite site hypothesis is still larger for the exome data of the myeloproliferative neoplasm ^31^. For patient 5 of the leukemia panel data ^34^ the BF is large enough that the posterior odds are certainly in favor of the infinite sites hypothesis being violated. These data are then the ‘smoking gun’ showing that the possibility of infinite sites violations needs to be seriously considered and treated for single-cell data. Again larger sample sizes will be needed to better assess the practical implications of these findings but modeling single cell data while allowing violations of the infinite sites hypothesis provides the statistical framework for exactly that.

The possibility of violations of the infinite sites assumption necessitates substantial adaptations in present-day models for reconstructing mutation histories of tumors. For example, in models designed for bulk sequencing data, a core assumption to deconvolve admixed mutation profiles is that the cellular frequency of a point mutation distributes over a single clade in the tumor phylogeny, a restriction that is contrary to the recurrence of a mutation in different parts of the tree. When looking at models based on single-cell data such as SCITE ^18^, the changes necessary to accommodate finite sites seem less profound, as indicated by the extension introduced in this paper to allow a single recurrent mutation. We also employed this method to search for multiple recurrences by restricting the recurrence to parallel mutations in data where higher scoring back mutations had been observed. This uncovered evidence of multiple violations of the ISA, but a strict statistical test would need to account for the higher scoring recurrences as well. However the generalization towards the recurrence of an unknown number of mutations in unknown multiplicities entails a vast extension of the underlying search space.

For single-cell data, we additionally have the issue of high doublet rates which, as we have seen, can severely affect reconstruction quality when not being explicitly modeled. While the accidental sequencing of more than one cell could be relatively easily prevented by rigorously checking samples prior to sequencing, it is likely to take some time before this issue is solved reasonably well for all technology platforms including high-throughput assays. Meanwhile it is essential to integrate doublets in models for reconstructing mutation histories from single-cell data. Especially for testing the ISA, modeling doublets is necessary since even a small number of doublets can interfere with the test. As we have shown in this work, modeling doublets is straightforward for a mutation-centric approach like SCITE ^18^. For sample-centric approaches such as BitPhylogeny ^16^ and OncoNEM ^17^, the integration of doublets may be a bit more involved, as the topology underlying the evolutionary history is no longer tree-like in the presence of admixed samples.

We focused in this work on testing the infinite sites assumption for point mutations in tumor evolution. This extends more generally to any cell lineages and their phylogeny where we know that violations become increasingly likely for larger sets of cells and mutations. Looking at larger-scale lesions in cancer, such as copy number alterations, the importance of allowing recurrent mutations becomes even more pronounced. These alterations typically affect larger segments which make it much more likely that the same site is affected multiple times. To model this type of lesions either alone or together with SNVs to integrate LOH, dropping the infinite sites assumption becomes even more crucial. Recent work using the less restrictive infinite alleles assumption ^21^ or Dollo parsimony ^22^ are promising first steps, but additional work on accurate models of tumor evolution and their inference from data is essential.

## Acknowledgements

We thank Giusi Moffa for very useful discussions about the Bayes Factor comparison ^29^ and Jochen Singer for bioinformatics support with the leukemia data^34^.

## Author contributions

JK and KJ developed and implemented the method. All authors conceived and designed the study. All authors drafted the manuscript and approved the final version.

## Correspondence

Correspondence and requests for materials should be addressed to Niko Beerenwinkel.

## Supplementary Information

### Methods

#### Tree models

The genealogy of somatic cells can be represented as a cell lineage tree, a rooted labeled binary tree, where the leaves represent the cells and the tree structure reflects the cell division history. Tree edges are labelled with mutation events and all cells below a mutation can be expected to exhibit this mutation. (See the left-most tree in Figure M1(a) for an example).

**Figure M1:**
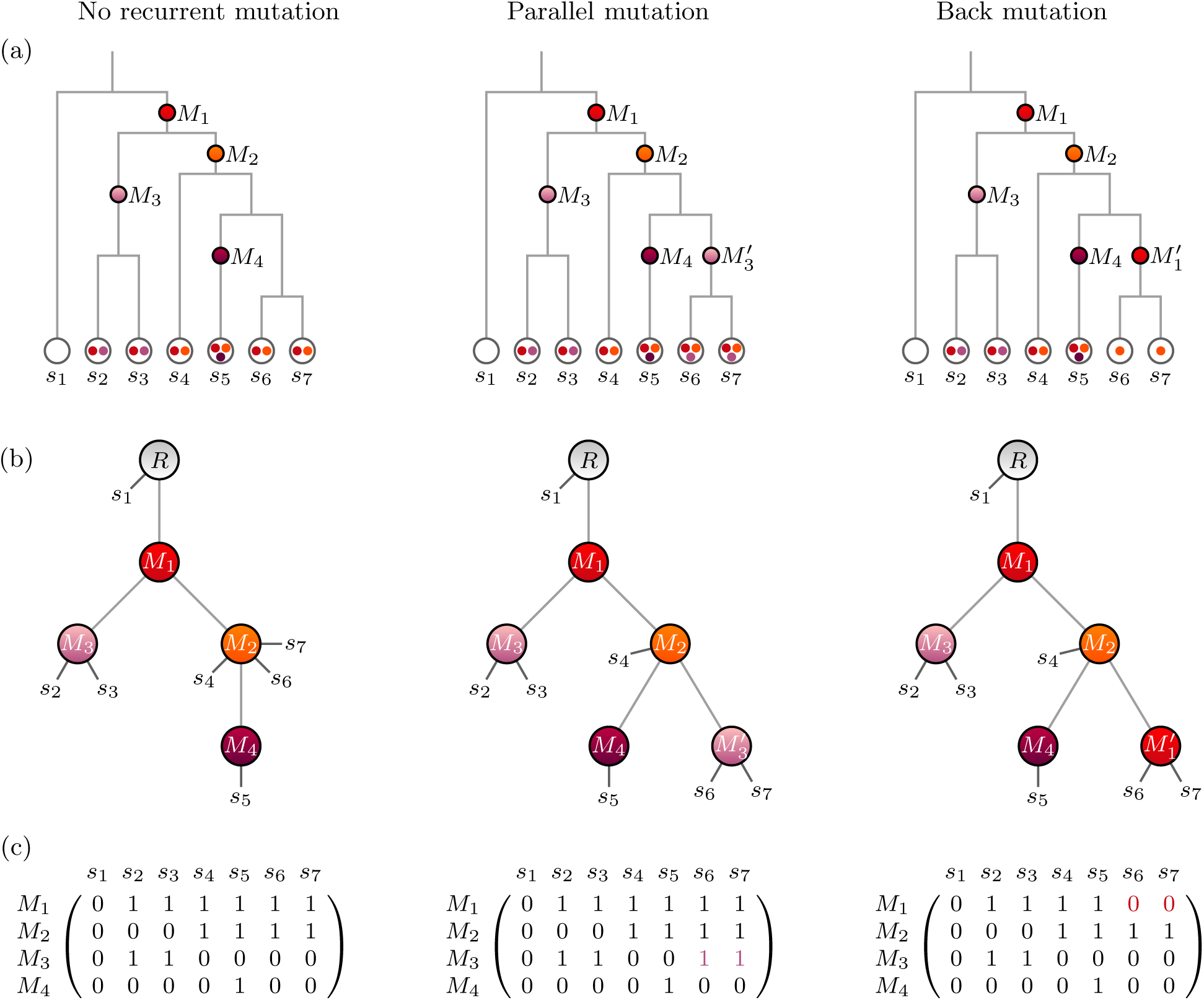
(*a*) Cell lineage trees of seven cells. *Left:* no recurrent mutations; *middle:* parallel mutation, a mutation occurs twice in separate lineages, denoted as *M_3_* and 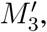, cells below both occurrences exhibit this mutation; *right*: back mutation, a second occurrence of a mutation in the same lineage brings the genomic site back to the original state, *i.e.* cells located below 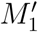 do not exhibit this mutation; (*b*) Mutation trees with attached cell samples. Each tree corresponds to the cell lineage tree in the same column; (*c*) Mutation matrices with binary states, each corresponds to the mutation tree in the same column; entry (*i, j*) contains the expected state of mutation *M_i_* in cell *S_j_*, 0 for absence and 1 for presence in the cell. The highlighted zeros in the matrix on the right are due to the placement of cells *s*_6_ and *s*_7_ below 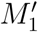, the second occurrence of mutation *M*_1_, which brings the genomic site back to the original state.

Models for somatic cell evolution typically make the infinite sites assumption which restricts any genomic site to host no more than one mutation event. Dropping this assumption means allowing not just one but multiple occurrences of the mutations in a cell lineage tree. For simplicity we allow here just a single mutation to occur twice. If the two copies of the same mutation happen in different branches, we refer to them as *parallel mutations.* A mutation that occurs twice in the same lineage represents a *back mutation.* We interpret this as the second mutation undoing the first mutation such that samples which have two copies of a mutation in their history would not exhibit the mutation. See Figure M1(a) for an illustration.

In SCITE ^18^ we utilized *mutation trees* as an alternative representation of mutation histories. The mutations form the tree nodes which are connected based on their partial temporal order (Figure M1(b)). A root is added to define the direction of the tree. Cell samples may attach to any of the nodes, and we expect them to contain all mutations on the path from the root to their attachment point. As with cell lineage trees, we can have parallel and back-mutations. The complete mutation history is defined by a pair (*𝑇*, ***σ***) where *𝑇* is the mutation tree and ***σ*** is the attachment array in which entry *j* encodes the node at which sample cell *S_j_* attaches to the mutation tree. For the trees in Figure M1 we have the attachment vectors

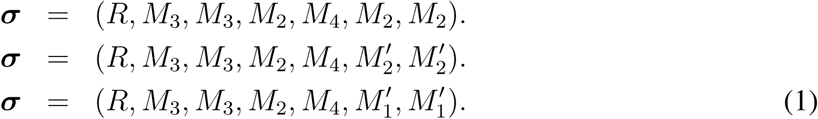

The mutation states of the cell samples can also be represented as a *mutation matrix E*. Here, entry (*i, j*) encodes the presence of a mutation *M_i_* in a cell *S_j_* with a 1 and its absence with a 0 (Figure M1(c)). In practice it is not necessary to construct the complete mutation matrix, as its entries can be obtained from *T* and ***σ****_j_*, the *j*-th entry of the attachment vector. Let anc*_T_*(***σ**_j_*) be the set of mutations that are ancestors of ***σ**_j_* in *T* including ***σ**_j_* itself, then we have

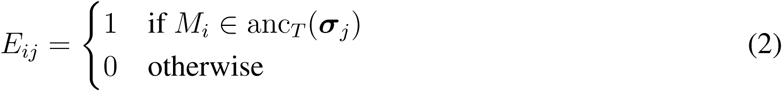

if *M_i_* is a unique mutation. For *M_i_* and 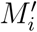 being the two incidences of a recurrent mutation, we have

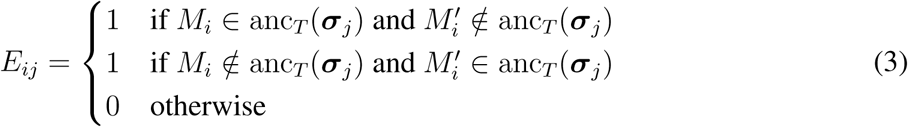

to encode the state after the mutation loss as a 0 in the mutation matrix.

#### Error model

In practice we observe a noisy version *D* of the expected mutation matrix *𝐸*. If the true mutation value is 0, we may observe a 1 with a probability of *α* (false positive) and if the true value is 1 we may observe a 0 with probability *β* (false negative):

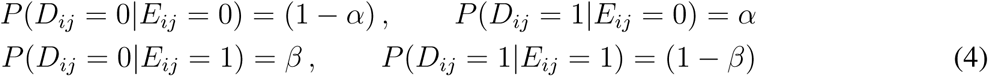

Assuming the observational errors are independent of each other, the likelihood of the data given a mutation tree *𝑇* and knowledge of the attachment of the samples *σ* is

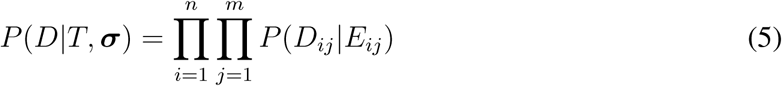

where *𝐸* is the expected mutation matrix for *𝑇* and ***σ*** To obtain the marginal tree likelihood independent of attachments, we sum over all attachment vectors ***σ***

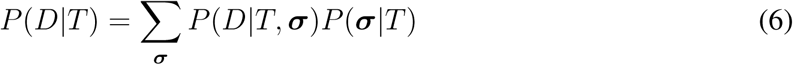

With *m* cells and *n* mutations, this can be computed efficiently in time *O*(*mn*) ^18^. Using a uniform prior for the sample attachment, *P*(***σ***|*𝑇*) becomes just a normalization constant that can be taken out of the sum. In the following we refer to the unnormalized marginal likelihood as the *tree score*

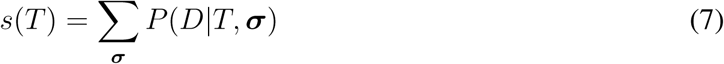

#### Modeling doublets

In single-cell sequencing it can happen that accidentally two (or more) cells are processed together which generates an admixed mutation profile of these cells (Figure M2). For our binary mutation states we assume that a mutation is called whenever it is present in at least one of the cells.

**Figure M2:**
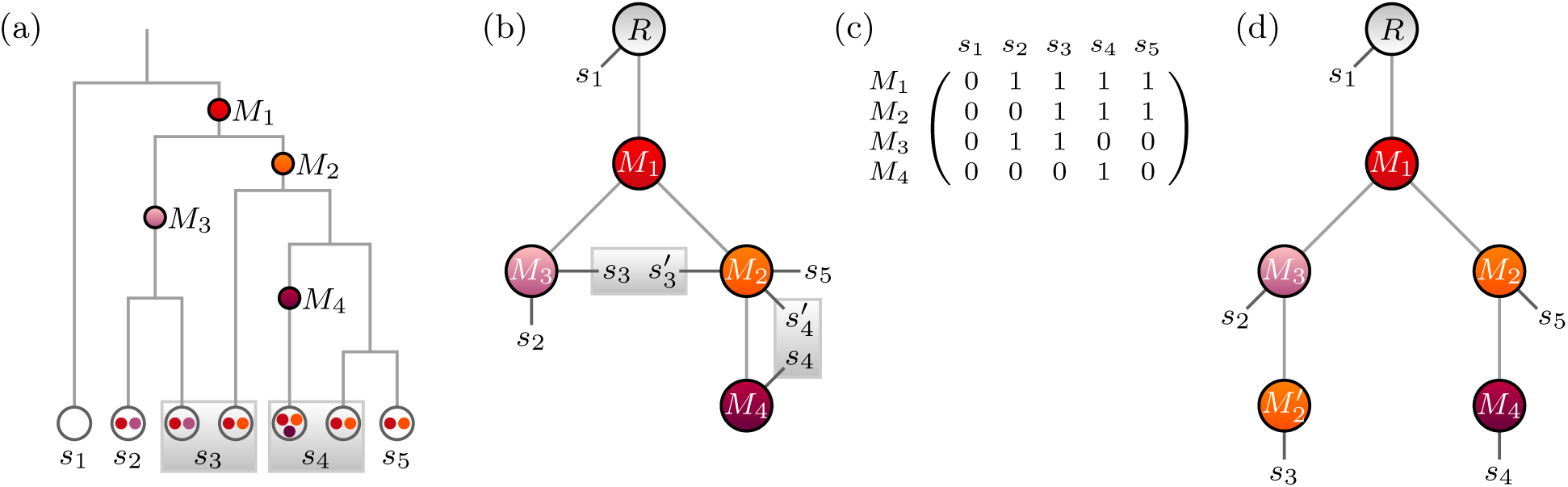
Tree reconstruction in the presence of doublet samples: (a) cell lineage tree with doublets (grey boxes). (b) Mutation tree with true sample attachment. Doublet samples (*s*_3_,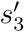) and (*s*_4_, 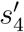) each attach to two different nodes. (c) mutation matrix with combined mutation states for the doublet samples. Mutation counted as present in a doublet sample, if present in at least one of the cells. (d) A tree with a recurrence of mutation *M*_2_ and no doublets is an alternative explanation for the mutation matrix in (c).

The two cells of a doublet sample *s_j_* can attach to different nodes of the mutation tree. Hence we change the attachment vector such that each entry *j* consists of a pair (***σ****_j_*, 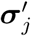) to indicate the two attachment points. The expected mutation vector is then defined as

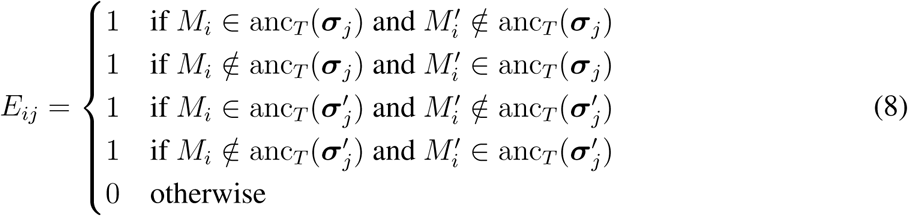

To accommodate for doublets in our model, we allow each sample to be a doublet with probability *δ* and a single cell with probability (1–*δ*). To obtain the tree likelihoods *Ṕ*(*𝐷*|*𝑇*) under this model, we first consider each sample separately. Let *𝐷_j_* be the observed mutation profile of sample *s_j_*, then we denote as

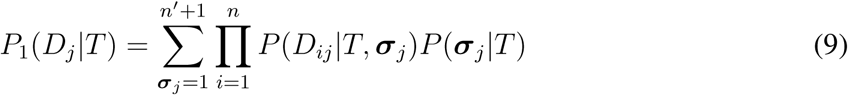

the likelihood of the tree for sample *s_j_* under the assumption that the sample is a single cell. The use of *n′* + 1 instead of *n* + 1 in the sum, accounts for the changing tree size when recurrences are allowed. For *n* mutations with a single recurrence, we have *n′* = *n* +1 tree nodes apart from the root, while *n′* = *n* in case of *n* unique mutations. Similarly we obtain

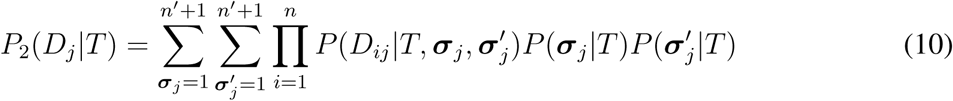

for the case that *s_j_* consists of two cells. To combine the two likelihoods we weight them by the respective single-cell and doublet probability.

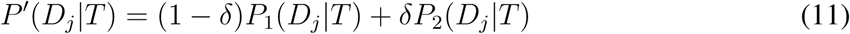

Then assuming that the sample attachments are independent of each other, the complete likelihood is the product over all samples:

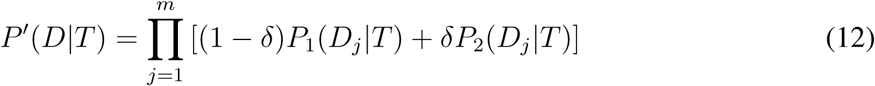

Since we have to account for all pairings of cells, the time complexity of calculating the likelihood is *O*(*m*^2^*n*). To obtain a tree score analogous to Equation (7), which is more useful for combinatorial considerations later, we divide the tree likelihood by the prior probability for a single attachment, a factor shared by all terms of the sum

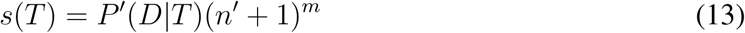

#### Model selection

To test the infinite sites hypothesis, we compare the evidence our observed data *𝐷* provides in favor of model *𝓜*_I_ consisting of trees with unique mutations and a model which allows for recurrent mutations. For simplicity we focus here on the model *𝓜*_F_ with exactly one repeated mutation. Finding strong evidence to favor *𝓜*_F_ over *𝓜*_I_ would be sufficient to reject the infinite sites hypothesis. We use Bayes factors for the model selection:

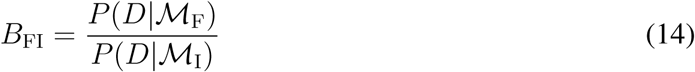

A value of *𝐵*_FI_ > 1 means that the data is better explained by the finite sites model than by the infinite sites model. The larger the number the stronger the evidence. To obtain the likelihood of *𝓜*_I_, we sum over all mutation trees with a single node for each mutated site observed in *𝐷* which gives us

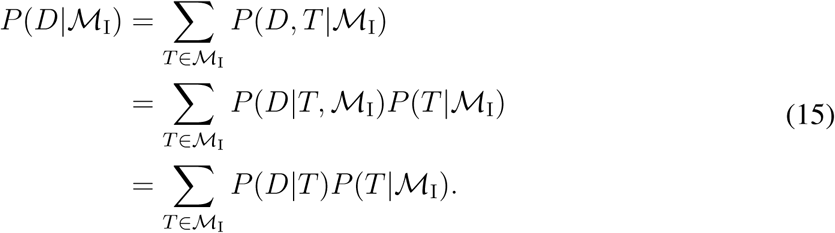

The dependency on *𝓜*_I_ in *𝑃*(*𝐷*|*𝑇*) can be dropped, as the data is no longer influenced by the model once the tree is fixed. To obtain the likelihood of a tree, we sum over all attachment vectors, such that

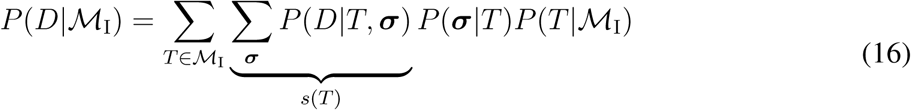

The unnormalized marginal tree likelihood is the tree score *s*(*𝑇*) as defined in Equation (7). Lastly using a uniform distribution for the prior on trees and sample attachments under a given model, we obtain

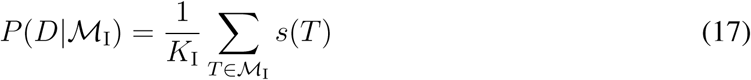

where *𝐾*_I_ is the number of pairs (*𝑇*, ***σ***) belonging to *𝓜*_I_,

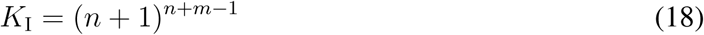

The finite sites model is the union of models *𝓜*_1_, …, *𝓜_n_*, where each *𝓜_i_* comprises all trees where only mutation *i* has a second occurrence. We then have

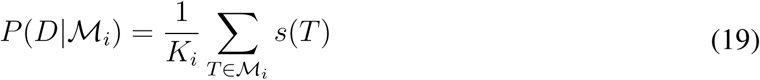

where *𝐾_i_* is the number of pairs (*𝑇*,***σ***) belonging to *𝑀_i_*.

However for the model comparison, we are only interested in trees that do not just recreate trees from the infinite sites model in the sense that the recurrent mutation does not give rise to any additional mutation profiles compared to a tree without the recurrence. For example if the recurrent mutation is the direct child or parent of the original copy, or if it shares a parent, then the recurrence can be removed. Excluding such trees from consideration and our model space leads to

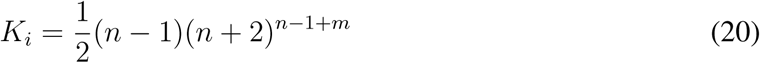

as derived in the Supplementary Material.

Another simple example is the case where the recurrent mutation has no descendant mutations and no samples attached to it. Then we recover a tree with no recurrent mutation and we should also exclude such cases. This possibility however depends on where the samples attach, rather than just on the tree so it cannot be excluded from the model space without interfering with the marginalization over ***σ***. Instead we include this possibility in our model class, along with further cases discussed in the Supplementary Material, but correct for their effect by deriving a lower bound for the number of trees and attachment pairs in *𝓜*_F_ that truly make use of the recurrent mutation and employ

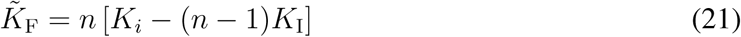

Likewise we obtain lower bounds for the model likelihood and the Bayes factor

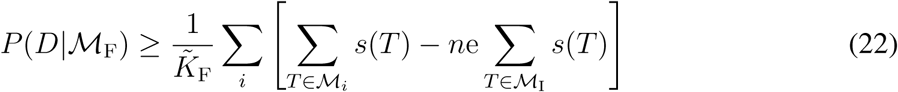

and

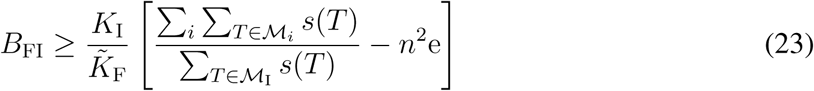

The derivation of the bounds is detailed in the Supplementary Material.

When calculating the tree scores, we take fixed values for the error rates, either those provided with the data or learnt under the infinite sites model. For the double rate *δ*, for each tree we find the value which maximize the score with numerical optimization equivalent to the EM algorithm. We also calculate as *δ̂* the fraction of samples which are doublets involving mutations from two lineages, since those doublets from a single lineage could be modeled as singlets instead.

#### Approximation

Typically there will be one recurrent mutation that increases the likelihood of the finite sites model much more strongly than the others

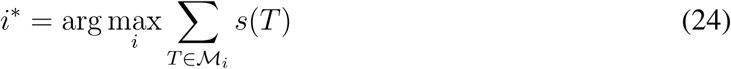

so that in the sum over *i* in Equation (23) the terms for the other *𝑀_i_* can essentially each be replaced by the sum over the copies of trees inside *𝓜*_I_ and

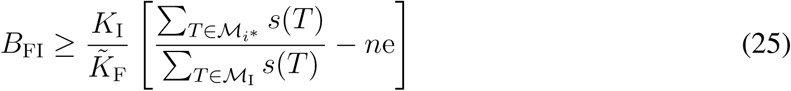

#### Estimation via MCMC

In general the sum over all tree scores can not be computed for the two models as both comprise a vast number of trees which grows super-exponentially in the number of mutations. Instead we estimate this value for each model using the MCMC scheme developed in SCITE ^18^ to search the space of rooted mutation trees.

Given the current tree *𝑇* we propose a tree *𝑇́* from the same model according to one of the three move types with some proposal probability *q*(*𝑇́*|*𝑇*) and accept the move with probability

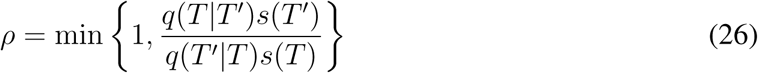

so that we obtain (after some burn in time) a sampler that provides trees proportionally to *s*(*𝑇*). Running the sampler for enough steps we will not only find a tree with the best score in a model *𝓜*,

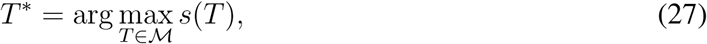

but also the total number of trees with the optimal score,

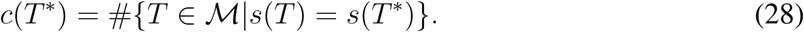

In general the optimal tree will be unique because of the marginalization over the attachment of samples (or we have two equivalent copies with labels swapped in *𝓜*_F_). If two or more mutations appear in exactly the same set of sampled cells (up to missing data) then the number of trees will grow, but equally for both model classes so the factors of *c* will anyway cancel. For completeness though we treat the arbitrary case here.

For our estimation of the sum score we now make use of the fact that in a sequence of trees sampled after burn-in, the fraction of optimal trees approximates the ratio between the sum score of all optimal trees and the sum score over the whole tree space:

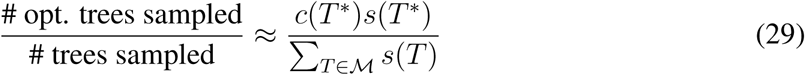

For this approximation to work, we need to know how long the MCMC needs to run until it has certainly converged. Then the chain of each run is equally likely to discover each of the maximally scoring trees. With the number of currently discovered maximal trees and the probability of discovering each of them in a run (or per state in the chain and the correlation between states) we can estimate and bound the probability that we are still missing any maximally scoring trees (or even better scoring ones). We can then simply run enough chains to reduce this to very low values.

For the left part of Equation (29) we simply need to know how often we hit a maximal tree in a typical chain. For this we run the chain several times and record, after a burn in period, the time the chain spends at the maximal score.

Simply running this procedure once for *𝓜* = *𝓜*_I_ and once for *𝓜* = *𝓜*_*i\**_ then allows us to find an approximation for the ratio in Equation (25). Running many chains gives confidence intervals on the ratios and hence on the final BFs.

#### Approximation via *s*(*𝑇*)

For some data the posterior may be very flat which prohibits sampling of the set of maximum scoring trees in reasonable time. For such cases, we make the approximation that the ratio of sampling the optimal trees in each model class is the same for both model classes so that

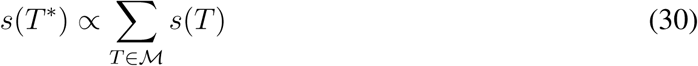

with the same proportionality constant for both model classes. Then we can effectively replace the sum over all trees in the BFs by *s*(*𝑇*^*^), the score of a single maximum scoring tree.

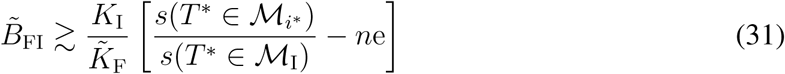

Finding the maximal score can also be made more efficient by monotonically changing the score landscape (for example raising the score to some power *γ*) which can be adapted to speed up the MCMC search.

